# In Vitro Activity of Eravacycline in Combination with Colistin Against Carbapenem-Resistant *A. baumannii* Isolates

**DOI:** 10.1101/552547

**Authors:** H. Selcuk Ozger, Tugba Cuhadar, Serap Suzuk Yildiz, Zehra Demirbas Gulmez, Murat Dizbay, Ozlem Guzel Tunccan, Ayşe Kalkanci, Husniye Simsek, Ozlem Unaldi

**Affiliations:** Department of Infectious Diseases and Clinical Microbiology, Gazi University School of Medicine, Ankara, Turkey; Department of Clinical Microbiology, Gazi University School of Medicine, Ankara, Turkey; MoH General Directorate of Public Health, Department of National AMR Surveillance Laboratory, Ankara, Turkey; MoH General Directorate of Public Health, Central Laboratory, Ankara, Turkey

**Author notes:** Address correspondence to H.Selcuk Ozger, Mevlana Street, No; 89, 06560, Ankara, Turkey.

## Abstract

The synergistic activity of eravacycline in combination with colistin on carbapenem-resistant *A.baumannii* (CRAB) isolates was evaluated in this study. Minimum inhibitory concentrations (MICs) of eravacycline and colistin were determined by the broth microdilution method. MICs values ranged between 1 to 4 mg and 0,5 to 128 mg/L for eravacycline and colistin, respectively. In-vitro synergy between eravacycline and colistin was evaluated by using the chequerboard methodology. Synergistic activity was found in 10 % of the strains, and additive effect in 20 %. No antagonism was detected. Similar activity was also observed in colistin resistant CRAB isolates. The result of this study indicates that eravacycline and colistin combination may be a potential therapeutic option for the treatment of CRAB related infections.

## Introduction

Carbapenem-resistant *Acinetobacter baumannii* (CRAB) is an important nosocomial pathogen causing substantial morbidity and mortality (1, 2). Current treatment options (polymyxins, tigecycline, aminoglycosides) for CRAB are limited and suffer from pharmacokinetic limitations such as high toxicity and low plasma levels (3). Therefore, to enhance clinical efficacy and to avoid toxicity, combination therapy is often used for CRAB infections (4).

Eravacycline is a novel fluorocycline that belongs to the tetracycline class of antimicrobials may be a treatment option for CRAB (5). In vitro studies were shown that eravacycline minimum inhibitory concentrations (MICs) were found to be 2-8 fold lower than tigecycline MICs against CRAB (6–8). The drug is also active against colistin resistant strains (9). Despite in vitro activity, treatment success of eravacycline in CRAB infections are unknown due to lack of clinical trials (10, 11). Management of healthcare-associated infections (HCAI) caused by CRAB should be required combination therapy. There is no *in vitro* study related to the synergism of eravacycline with other antibiotics in the literature. The aim of this *in vitro* study is to evaluate the synergistic activity of eravacycline in combination with colistin on CRAB isolates.

## Results

Ten carbapenem-resistant *A. baumannii* strains were used in this study. Three of these strains were also resistant to colistin. All isolates were found to have *bla*_OXA-51_ and nine harbored *bla*_OXA-23._ The characteristics of 10 CRAB isolates included in chequerboard experiments were shown in Table 1.

**Table 1.**
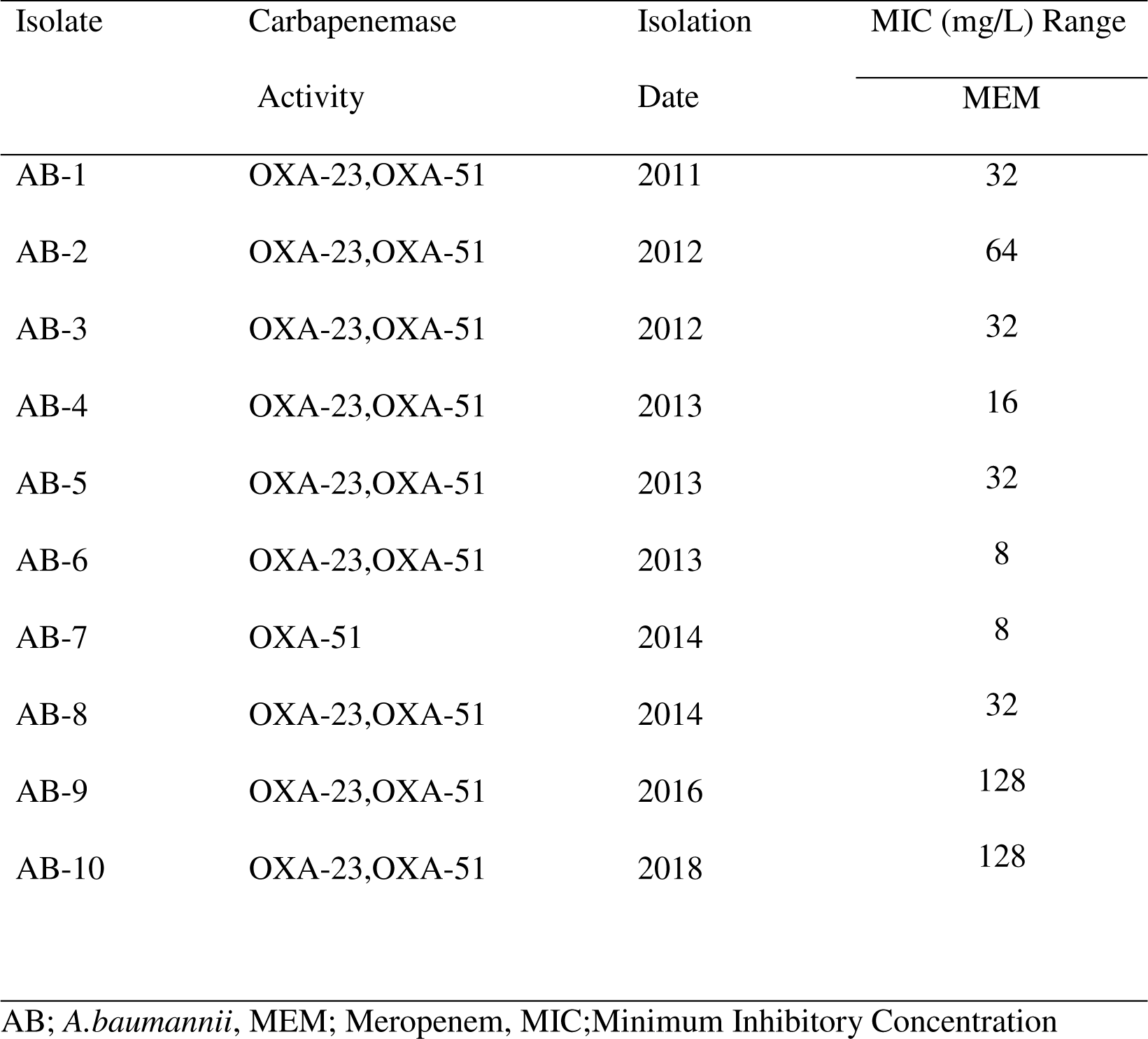
Characteristics of Carbapenem Resistant *A.baumannii* Isolates

| Isolate | Carbapenemase | Isolation | MIC (mg/L) Range |
| --- | --- | --- | --- |
|  | Activity | Date | MEM |
| AB-1 | OXA-23,OXA-51 | 2011 | 32 |
| AB-2 | OXA-23,OXA-51 | 2012 | 64 |
| AB-3 | OXA-23,OXA-51 | 2012 | 32 |
| AB-4 | OXA-23,OXA-51 | 2013 | 16 |
| AB-5 | OXA-23,OXA-51 | 2013 | 32 |
| AB-6 | OXA-23,OXA-51 | 2013 | 8 |
| AB-7 | OXA-51 | 2014 | 8 |
| AB-8 | OXA-23,OXA-51 | 2014 | 32 |
| AB-9 | OXA-23,OXA-51 | 2016 | 128 |
| AB-10 | OXA-23,OXA-51 | 2018 | 128 |

AB; *A.baumannii*, MEM; Meropenem, MIC; Minimum Inhibitory Concentration

MIC values ranged between 1 to 4 mg and 0,5 to 128 mg/L for eravacycline and colistin, respectively. Chequerboard analysis showed 10% synergy, 20% additive, 70% indifference. No antagonism was observed. In both colistin and carbapenem-resistant strains, one strain showed synergy, and two of them were indifferent. The MIC values of eravacycline-colistin and minimum FIC values are summarized in Table 2.

**Table 2.** Synergistic Activity of Eravacycline in Combination with Colistin

| Isolate | MIC (mg/L) Range | | $\Sigma$ FIC<sub>min</sub> | Interpretation |
| --- | --- | --- | --- | --- |
|  | Eravacycline | Colistin |  |  |
| AB-1 | 2 | 1 | 0,75 | Add |
| AB-2 | 1 | 1 | 1,03 | Ind |
| AB-3 | 2 | 0,5 | 1,06 | Ind |
| AB-4 | 2 | 1 | 1,03 | Ind |
| AB-5 | 1 | 1 | 1,06 | Ind |
| AB-6 | 2 | 32 | 1,00 | Ind |
| AB-7 | 4 | 1 | 1,03 | Ind |
| AB-8 | 4 | 128 | 0,50 | Syn |
| AB-9 | 2 | 256 | 1,01 | Ind |
| AB-10 | 4 | 0,5 | 0,53 | Add |

MIC, minimum inhibitory concentration ; $\Sigma$ FIC min, smallest total fractional inhibitory
concentration ; Ind, indifference ; Add, Additive; Syn, synergy

## Discussion

To our knowledge, this is the first study assessing the synergistic activity of eravacycline with colistin. Our study demonstrated that eravacycline had synergistic and additive activity in combination with colistin without antagonism. Our results revealed that the combination of eravacycline and colistin may be a treatment option for CRAB related infections, including colistin-resistant isolates.

*A. baumannii* is the most common nosocomial pathogen that causes bloodstream infections (BSI) or ventilator-associated pneumonia (VAP) in many medical centers, particularly in Turkey (2, 5). Carbapenem resistance rates exceed 90% in some parts of the World (3). Carbapenem resistance among *A. baumannii* isolates in Turkey is ranged from 91% to 98% according to national health statistics (12).

Therapy of CRAB infections is generally failing and limited with colistin, mostly (8). Because of increasing resistance rates and pharmacokinetic limitations, colistin often used in combination with other antibiotics such as meropenem, tigecycline, rifampicin. However, the optimum treatment regimen is still uncertain (1, 13–15). There is no doubt that new therapeutic options are urgently needed for the treatment of CRAB infections.

Eravacycline is a novel fluorocycline, with a tetracycline core, that binds to the 70S ribosome submit of bacteria (8, 16). The activity of eravacycline against gram-negative, gram-positive and anaerobic bacteria except *P.aeruginosa* was demonstrated in previous studies (7, 8, 17–19). Furthermore, eravacycline was active against multidrug-resistant bacteria, including those expressing carbapenemases (7). It was utilized as a potent antibiotic for *A.baumannii*, including isolates associated with an acquired OXA or up-regulation of the intrinsic OXA-51-like enzyme (1, 6, 9). The MIC 50/90 values were 0.5 and 1 mg/L and higher in tigecycline resistant strains (1, 6, 9). The MIC values did not change with the production of OXA carbapenemases (1). In accordance with the previous studies, eravacycline MIC values ranged from 0,5-4 mg/L for OXA-23 and OXA-51 producing CRAB strains in our study. Considering the prevalence of OXA-type carbapenemase activity in CRAB strains, eravacycline has been evaluated as a good treatment option (20–22). However, some type of carbapenemase activity, increased expression of the efflux pumps and the presence of tigecycline resistance are associated with increasing MIC values for eravacycline (1, 3).

Therefore, to restrict the emergence of resistance, combination therapies include eravacycline might be an option for treating CRAB infections (4). Colistin is the most commonly used antibiotic in combination therapies in HCAI caused by CRAB (4, 23, 24). However, colistin resistance is gradually increasing especially in CRAB strains. Colistin-resistant CRAB isolates were also included in our study. We found synergy (in one strain) and indifference (in two strain) in colistin resistant strains. This preliminary in vitro results clarified that eravacycline and colistin combination should be evaluated with randomized clinical studies.

Synergistic activities of antimicrobial agents have studied by chequerboard microdilution method and simultaneous time-kill analysis. Synergy frequencies may vary depending on the method used. Compared with the time kill method, the chequerboard method reveals lower synergy frequencies (4). In our study, the in-vitro synergy between eravacycline and colistin was evaluated by using the chequerboard methodology. Therefore, we have limitations in our study. Our results should be evaluated and supported by other in vitro methods. Moreover, in vitro studies can provide preliminary guidance for rational drug combination use in the clinics.

In conclusion, we found 10% synergy and 20% additive activity without antagonism between eravacycline and colistin in CRAB isolates. Similar activity is maintained in colistin resistant CRAB isolates. The result of this study indicates that eravacycline and colistin combination may be a potential therapeutic option for the treatment of CRAB related infections.

## Acknowledgments

This research received no specific grant from any funding agency in the public, commercial, or not-for-profit sectors.

## Material-Methods

### I. Bacterial Strains

Carbapenem-resistant *A. baumannii* isolates were used in the study. All strains were identified in lower respiratory tract samples in critically ill patients. Identification on species levels was determined by MALDI-TOF MS (Bruker Biotyper; Bruker Daltonics, Bremen, Germany). During testing, the isolates were cultured from frozen stocks with 5% sheep blood agar in accordance with guidelines from the Clinical and Laboratory Standards Institute (25). All strains were incubated 35°C before testing.

### II.Investigation of Carbapenemase Resistance

Minimum inhibitory concentrations of meropenem was determined for all strains by the broth microdilution method (25). MIC values of 4 µg/ml and above are taken as limit values for meropenem resistance (26). All strains screened for carbapenemase activity by polymerase chain reaction (PCR). Eight of the most common carbapenemase genes (*bla*_OXA-23_, *bla*_OXA-48_, *bla*_OXA-51_, *bla*_OXA-58_, *bla* _NDM_, *bla*_IMP_, *bla*_VIM_, and *bla*_KPC_) were screened by an in-house multiplex PCR test (27-33). The oligos used for the amplification of the genes as shown in Table 3.

**Table 3.**
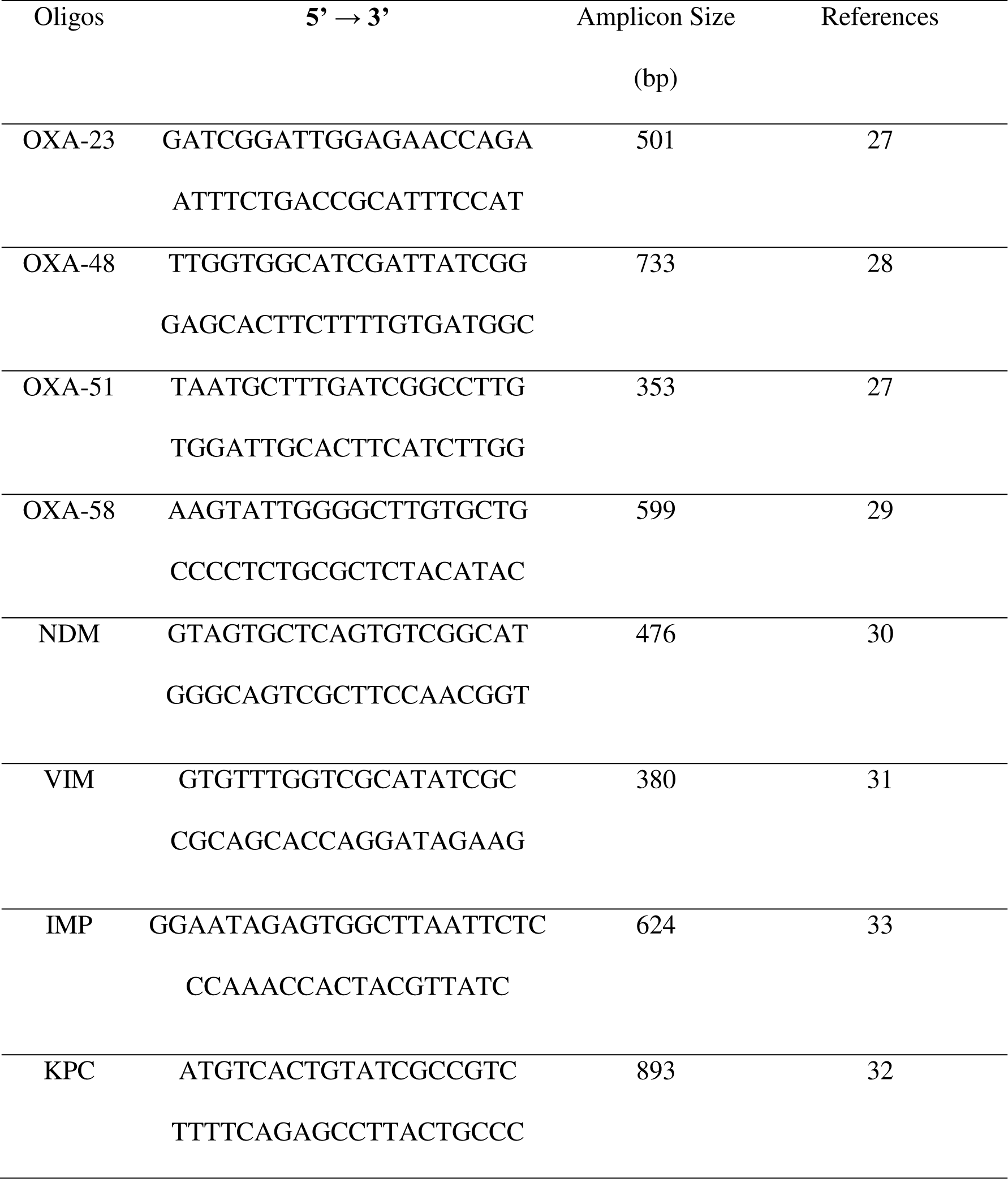
Oligos Used for Amplification

### III.Drugs

Eravacycline (Lot number: 26030) and colistin (Lot number:16647) were provided MedChemTronica (Sweden) as laboratory-grade powders. All drugs were dissolved with dHO_2_, 6,4 mg/ml stock solutions were prepared for eravacycline and colistin. All stock solutions were stored at – 20 °C throughout the study.

### IV.Minimum Inhibitory Concentration And Fractional Inhibitory Concentrations (FIC)

Minimum inhibitory concentrations of eravacycline and colistin were determined for all strains by the broth microdilution method (26). The data obtained from broth microdilution tests were used to calculate synergy. The chequerboard microdilution panel method was used for MIC determination of the eravacycline/colistin combination.

Using 96-well U-bottom microplates, graded concentrations of antibiotics were mixed. Each antimicrobial agent was prepared to a fixed volume of 50 µL (up to a total of 100 µL volume for two antimicrobial agents), and 10 µL of bacterial suspension was added to each well. The final concentration of the test strains was a 5×10 ^5^ CFU/mL in a total final volume of 100 µL in each well. The plates were incubated for 16-24h at 35°C and the presence or inhibition of microbial growth was determined visually. Eravacycline and colistin synergy were studied at least 2 times with the chequerboard method in all strains. The fractional inhibitory concentration (FIC) index was calculated with the formula:

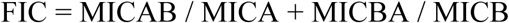

The results of combination tests according to the FIC index were interpreted as follows: synergistic (FIC ≤ 0.5), additive (FIC >0.5 and ≤1), indifferent (FIC >1 and ≤4) and antagonistic (FIC >4).

## References

1. Paul M, Daikos GL, Durante-Mangoni E, Yahav D, Carmeli Y, Benattar YD, Skiada A, Andini R, Eliakim-Raz N, Nutman A, Zusman O, Antoniadou A, Pafundi PC, Adler A, Dickstein Y, Pavleas I, Zampino R, Daitch V, Bitterman R, Zayyad H, Koppel F, Levi I, Babich T, Friberg LE, Mouton JW, Theuretzbacher U, Leibovici L. 2018. Colistin alone versus colistin plus meropenem for treatment of severe infections caused by carbapenem-resistant Gram-negative bacteria: an open-label, randomised controlled trial. Lancet Infect Dis 18:391–400.

2. Aydin M, Ergonul O, Azap A, Bilgin H, Aydin G, Cavus SA, Demiroglu YZ, Aliskan HE, Memikoglu O, Menekse S, Kaya S, Demir NA, Karaoglan I, Basaran S, Hatipoglu C, Erdinc S, Yilmaz E, Tumturk A, Tezer Y, Demirkaya H, Cakar SE, Keske S, Tekin S, Yardimci C, Karakoc C, Ergen P, Azap O, Mulazimoglu L, Ural O, Can F, Akalin H. 2018. Rapid emergence of colistin resistance and its impact on fatality among healthcare-associated infections. J Hosp Infect 98:260–263.

3. Isler B, Doi Y, Bonomo RA, Paterson DL. 2019. New Treatment Options against Carbapenem-Resistant Acinetobacter baumannii Infections. Antimicrob Agents Chemother 63.

4. Ni W, Shao X, Di X, Cui J, Wang R, Liu Y. 2015. In vitro synergy of polymyxins with other antibiotics for Acinetobacter baumannii: a systematic review and meta-analysis. Int J Antimicrob Agents 45:8–18.

5. But A, Yetkin MA, Kanyilmaz D, Aslaner H, Bastug A, Aypak A, Onguru P, Akinci E, Mutlu NM, Bodur H. 2017. Analysis of epidemiology and risk factors for mortality in ventilator-associated pneumonia attacks in intensive care unit patients. Turk J Med Sci 47:812–816.

6. Livermore DM, Mushtaq S, Warner M, Woodford N. 2016. In Vitro Activity of Eravacycline against Carbapenem-Resistant Enterobacteriaceae and Acinetobacter baumannii. Antimicrob Agents Chemother 60:3840–4.

7. Sutcliffe JA, O'Brien W, Fyfe C, Grossman TH. 2013. Antibacterial activity of eravacycline (TP-434), a novel fluorocycline, against hospital and community pathogens. Antimicrob Agents Chemother 57:5548–58.

8. Abdallah M, Olafisoye O, Cortes C, Urban C, Landman D, Quale J. 2015. Activity of eravacycline against Enterobacteriaceae and Acinetobacter baumannii, including multidrug-resistant isolates, from New York City. Antimicrob Agents Chemother 59:1802–5.

9. Seifert H, Stefanik D, Sutcliffe JA, Higgins PG. 2018. In-vitro activity of the novel fluorocycline eravacycline against carbapenem non-susceptible Acinetobacter baumannii. Int J Antimicrob Agents 51:62–64.

10. Solomkin J, Evans D, Slepavicius A, Lee P, Marsh A, Tsai L, Sutcliffe JA, Horn P. 2017. Assessing the Efficacy and Safety of Eravacycline vs Ertapenem in Complicated Intra-abdominal Infections in the Investigating Gram-Negative Infections Treated With Eravacycline (IGNITE 1) Trial: A Randomized Clinical Trial. JAMA Surg 152:224–232.

11. Solomkin JS, Gardovskis J, Lawrence K, Montravers P, Sway A, Evans D, Tsai L. 2018. IGNITE4: Results of a Phase 3, Randomized, Multicenter, Prospective Trial of Eravacycline vs. Meropenem in the Treatment of Complicated Intra-Abdominal Infections. Clin Infect Dis doi:10.1093/cid/ciy1029.

12. National Antimicrobial Reistance Surveillance Report, 2017, Turkey.((https://hsgm.saglik.gov.tr/tr/duyurular/997-ulusal-sağlık-hizmeti-ilişk-enfeksiyonlar-surveyans-ağı-etken-dağılımı-ve-antibiyotik-direnç-raporu-2017.html)

13. Amat T, Gutierrez-Pizarraya A, Machuca I, Gracia-Ahufinger I, Perez-Nadales E, Torre-Gimenez A, Garnacho-Montero J, Cisneros JM, Torre-Cisneros J. 2018. The combined use of tigecycline with high-dose colistin might not be associated with higher survival in critically ill patients with bacteraemia due to carbapenem-resistant Acinetobacter baumannii. Clin Microbiol Infect 24:630–634.

14. Vardakas KZ, Mavroudis AD, Georgiou M, Falagas ME. 2018. Intravenous colistin combination antimicrobial treatment vs. monotherapy: a systematic review and meta-analysis. Int J Antimicrob Agents 51:535–547.

15. Liang CA, Lin YC, Lu PL, Chen HC, Chang HL, Sheu CC. 2018. Antibiotic strategies and clinical outcomes in critically ill patients with pneumonia caused by carbapenem-resistant Acinetobacter baumannii. Clin Microbiol Infect 24:908 e1-908 e7.

16. Grossman TH, Starosta AL, Fyfe C, O'Brien W, Rothstein DM, Mikolajka A, Wilson DN, Sutcliffe JA. 2012. Target- and resistance-based mechanistic studies with TP-434, a novel fluorocycline antibiotic. Antimicrob Agents Chemother 56:2559–64.

17. Zhanel GG, Baxter MR, Adam HJ, Sutcliffe J, Karlowsky JA. 2018. In vitro activity of eravacycline against 2213 Gram-negative and 2424 Gram-positive bacterial pathogens isolated in Canadian hospital laboratories: CANWARD surveillance study 2014-2015. Diagn Microbiol Infect Dis 91:55–62.

18. Monogue ML, Thabit AK, Hamada Y, Nicolau DP. 2016. Antibacterial Efficacy of Eravacycline In Vivo against Gram-Positive and Gram-Negative Organisms. Antimicrob Agents Chemother 60:5001–5.

19. Zhanel GG, Cheung D, Adam H, Zelenitsky S, Golden A, Schweizer F, Gorityala B, Lagace-Wiens PR, Walkty A, Gin AS, Hoban DJ, Karlowsky JA. 2016. Review of Eravacycline, a Novel Fluorocycline Antibacterial Agent. Drugs 76:567–88.

20. Beris FS, Budak EE, Gulek D, Uzun A, Cizmeci Z, Mengeloglu FZ, Direkel S, Cetinkol Y, Ay Altintop Y, Iraz M, Dal T, Say Coskun SU, Balci PO, Kayman T, Caliskan A, Yazici Y, Tosun I, Erturk A, Copur Cicek A. 2016. [Investigation of the frequency and distribution of beta-lactamase genes in the clinical isolates of Acinetobacter baumannii collected from different regions of Turkey: a multicenter study]. Mikrobiyol Bul 50:511–521.

21. Pournaras S, Dafopoulou K, Del Franco M, Zarkotou O, Dimitroulia E, Protonotariou E, Poulou A, Zarrilli R, Tsakris A. 2017. Predominance of international clone 2 OXA-23-producing-Acinetobacter baumannii clinical isolates in Greece, 2015: results of a nationwide study. Int J Antimicrob Agents 49:749–753.

22. Nowak J, Zander E, Stefanik D, Higgins PG, Roca I, Vila J, McConnell MJ, Cisneros JM, Seifert H. 2017. High incidence of pandrug-resistant Acinetobacter baumannii isolates collected from patients with ventilator-associated pneumonia in Greece, Italy and Spain as part of the MagicBullet clinical trial. J Antimicrob Chemother 72:3277–3282.

23. Wang J, Niu H, Wang R, Cai Y. 2018. Safety and efficacy of colistin alone or in combination in adults with Acinetobacter baumannii infection: a systematic review and meta-analysis. Int J Antimicrob Agents doi:10.1016/j.ijantimicag.2018.10.020.

24. Mohammadi M, Khayat H, Sayehmiri K, Soroush S, Sayehmiri F, Delfani S, Bogdanovic L, Taherikalani M. 2017. Synergistic Effect of Colistin and Rifampin Against Multidrug Resistant Acinetobacter baumannii: A Systematic Review and Meta-Analysis. Open Microbiol J 11:63–71.

25. CLSI. 2012. Methods for Dilution Antimicrobial Susceptibility Tests f or Bacteria That Grow Aerobically; Approved St andard—Ninth Edition. CLSI document M07-A9. Wayne, PA: Clinical and Laboratory Standards Institute.

26. CLSI. 2018. Performance Standards for Antimicrobial Susceptibility Testing. 28th ed. CLSI supplement M100. Wayne, PA: Clinical and Laboratory Standards Institute.

27. Hou C, Yang F. 2015. Drug-resistant gene of blaOXA-23, blaOXA-24, blaOXA-51 and blaOXA-58 in Acinetobacter baumannii. Int J Clin Exp Med 8:13859–63.

28. Poirel L, Bonnin RA, Nordmann P. 2012. Genetic Features of the Widespread Plasmid Coding for the Carbapenemase OXA-48. Antimicrobial Agents and Chemotherapy 56:559–562.

29. Zhou H, Pi BR, Yang Q, Yu YS, Chen YG, Li LJ, Zheng SS. 2007. Dissemination of imipenem-resistant Acinetobacter baumannii strains carrying the ISAba1 blaOXA-23 genes in a Chinese hospital. J Med Microbiol 56:1076–80.

30. Mushtaq S, Irfan S, Sarma JB, Doumith M, Pike R, Pitout J, Livermore DM, Woodford N. 2011. Phylogenetic diversity of Escherichia coli strains producing NDM-type carbapenemases. J Antimicrob Chemother 66:2002–5.

31. Garza-Ramos U, Morfin-Otero R, Sader HS, Jones RN, Hernandez E, Rodriguez-Noriega E, Sanchez A, Carrillo B, Esparza-Ahumada S, Silva-Sanchez J. 2008. Metallo-beta-lactamase gene bla(IMP-15) in a class 1 integron, In95, from Pseudomonas aeruginosa clinical isolates from a hospital in Mexico. Antimicrob Agents Chemother 52:2943–6.

32. Gomez-Gil MR, Pano-Pardo JR, Romero-Gomez MP, Gasior M, Lorenzo M, Quiles I, Mingorance J. 2010. Detection of KPC-2-producing Citrobacter freundii isolates in Spain. J Antimicrob Chemother 65:2695–7.

33. Kaczmarek FM, Dib-Hajj F, Shang W, Gootz TD. 2006. High-level carbapenem resistance in a Klebsiella pneumoniae clinical isolate is due to the combination of bla(ACT-1) beta-lactamase production, porin OmpK35/36 insertional inactivation, and down-regulation of the phosphate transport porin phoe. Antimicrob Agents Chemother 50:3396–406.

